# Genomic prediction of metabolic content in rice grain in response to warmer night conditions

**DOI:** 10.1101/2024.07.23.604827

**Authors:** Ye Bi, Harkamal Walia, Toshihiro Obata, Gota Morota

## Abstract

It has been argued that metabolic content can be used as a selection marker to accelerate crop improvement because metabolic profiles in crops are often under genetic control. Evaluating the role of genetics in metabolic variation is a long-standing challenge. Rice, one of the world’s most important staple crops, is known to be sensitive to recent increases in nighttime temperatures. Quantification of metabolic levels can help measure rice responses to high night temperature (HNT) stress. However, the extent of metabolic variation that can be explained by regression on whole-genome molecular markers remains to be evaluated. In the current study, we generated metabolic profiles for mature grains from a subset of rice diversity panel accessions grown under optimal and HNT conditions. Metabolite accumulation was low to moderately heritable, and genomic prediction accuracies of metabolite accumulation were within the expected upper limit set by their genomic heritability estimates. Genomic heritability estimates were slightly higher in the control group than in the HNT group. Genomic correlation estimates for the same metabolite accumulation between the control and HNT conditions indicated the presence of genotype-by-environment interactions. Reproducing kernel Hilbert spaces regression and image-based deep learning improved prediction accuracy, suggesting that some metabolite levels are under non-additive genetic control. Joint analysis of multiple metabolite accumulation simultaneously was effective in improving prediction accuracy by exploiting correlations among metabolites. The current study serves as an important first step in evaluating the cumulative effect of markers in influencing metabolic variation under control and HNT conditions.

**Core ideas:**

- Rice is sensitive to increases in nighttime and daytime temperatures
- Metabolite accumulation from rice grains was low to moderately heritable
- Non-additive genomic prediction models improved prediction accuracy for some metabolites
- Results shed new light on the utility of genomic predictions for metabolite accumulation from rice grains

## Introduction

It has long been recognized that metabolic profiles in plants are under genetic control (Saito and Matsuda, 2010; Saito, 2013). It has also been argued that the use of metabolite accumulation as quantitative traits can be an important addition to the resources that can be used to accelerate crop improvement through genome-assisted breeding (Fernie and Schauer, 2009; Kliebenstein, 2009). Plant metabolites play essential roles in growth, development, agronomic performance, quality, flavor, and stress resistance. In particular, metabolites are considered intermediates of biochemical reactions that link agronomic traits to the genome. Thus, metabolite profiles provide an opportunity to understand the physiological state of a crop. Due to recent advances in metabolite profiling techniques, the combination of genetic and metabolic approaches is increasingly being used to study end-use quality traits in crop plants (Fernie and Tohge, 2017).

Rice (*Oryza sativa L.*) is one of the most important staple crops worldwide, providing energy and nutrients to meet the needs of a growing population. Several studies have characterized the metabolic profiling of diverse rice accessions and mature grain metabolites are considered important for yield and grain quality (Oikawa et al., 2008; Fitzgerald et al., 2009). Seed quality of rice is of agronomic importance, which is influenced by the chemical composition (Kusano et al., 2015). Rice produces many kinds of bioactive compounds. Previous studies have characterized metabolic differences between accessions or subpopulations (Kusano et al., 2007; Kim et al., 2013; Hu et al., 2014). One strategy for integrating metabolite accumulation in rice into quantitative genetic modeling is to treat metabolite accumulation as predictors or auxiliary phenotypes to improve the prediction accuracy of target traits (Bi et al., 2023). Genotype-metabolite association studies in rice have also been reported for primary and secondary (i.e., specialized) metabolites using QTL mapping (Matsuda et al., 2012; Ying et al., 2012; Gong et al., 2013) and genome-wide association studies (Chen et al., 2014; Matsuda et al., 2015; Chen et al., 2016) to dissect the biochemical basis of the rice metabolome. However, the investigation of the extent of metabolic variation that can be ex-plained by whole-genome molecular markers (e.g., genomic prediction) remains fragmentary. The availability of high-coverage metabolic and single nucleotide polymorphism (SNP) data provides an opportunity to perform quantitative genetic analysis of rice metabolite accumulation, such as estimation of genomic heritability and genomic correlation, or prediction of genetic values using whole-genome regression analysis (Meuwissen et al., 2001; Yang et al., 2010).

Although sustained increases in rice production and quality improvement are essential to meet the demands of a growing population, rice is susceptible to abiotic stresses such as heat, cold, drought, flooding, and salinity that result in yield losses (Wheeler and Von Braun, 2013; Zhao et al., 2017). Of these, a greater increase in nighttime temperatures than daytime temperatures is of particular concern (Vose et al., 2005; Donat and Alexander, 2012; Xia et al., 2014). Previous studies have reported that high night temperatures (HNT) negatively affect grain yield and quality traits (Morita et al., 2005; Welch et al., 2010; Peng et al., 2013; Sreenivasulu et al., 2015; Jagadish et al., 2015; Wang et al., 2017; Wada et al., 2019; Impa et al., 2021; Dhatt et al., 2021; Sakai et al., 2022; Sandhu et al., 2024). Because changes in biochemical or physiological levels under abiotic stress are likely to be reflected in the metabolic profiles of plants (Obata and Fernie, 2012), quantification of metabolic levels can help measure rice responses to HNT stress. A recent study showed differential metabolic abundance in rice grains between control and HNT conditions (Dhatt et al., 2019).

With the increasing impact of climate change on crop productivity, evaluating the role of genetic variation in regulating the abundance of rice metabolites under control and HNT conditions can be useful for understanding rice responses to HNT stress. Therefore, the objectives of this study were 1) to quantify the extent of variation in rice metabolite content explained by all SNP and 2) to evaluate the accuracy of genomic prediction of metabolite accumulation under control and HNT conditions.

## Materials and Methods

### Plant materials and growth conditions

Rice diversity panel 1 lines (Zhao et al., 2011) were used in this study. These lines (accessions) represent the following subpopulation groups: tropical japonica (25.11%), temperate japonica (22.37%), indica (18.72%), aus (17.35%), admixed japonica (9.13%), aromatic (3.20%), admixed indica (2.74%), and admixed (1.38%). Six seedlings per accession were transplanted into 4-inch pots containing natural soil. The pots were arranged in a randomized complete block design. The HNT experiment was conducted as previously described (Dhatt et al., 2021; Bi et al., 2023). Briefly, all plants were grown under controlled conditions until flowering. When approximately 50% of the primary panicle had completed fertilization, half of the plants from each accession were transferred to HNT conditions until maturity. All plants were harvested at physiological maturity. A total of 192 and 188 rice lines were obtained from the control and HNT conditions, respectively. All lines were genotyped using a highdensity rice array of 700k SNP markers (McCouch et al., 2016). A total of 388,219 SNP markers were used for analysis after removing SNP markers with minor allele frequencies less than 0.05.

### Metabolic profiling

The metabolic profiling procedures based on gas chromatography-mass spectrometry are described elsewhere (Bi et al., 2023). Briefly, five dehusked mature grains of each genotype were taken from the pool of all plant individuals and used for metabolite profiling. Grains were frozen and ground to a fine powder using a ball mill (TissuelyzerII, Qiagen, Düsseldorf, Germany) at liquid nitrogen temperature. An aliquot of approximately 50 mg was weighed and used for metabolite extraction and profiling using a 7200 GC-QTOF system (Agilent, Santa Clara, CA, USA) according to the previously described protocol (Wase et al., 2022). Chromatography peaks were annotated to metabolites according to retention time and mass spectral information in the Fiehn Metabolomics database (Agilent). Peak heights of representative ions for individual metabolites were normalized to that of the internal standard, ribitol (m/z 319), and to the fresh weights of the materials to determine relative metabolite content. Retention time and representative ion m/z of each peak and relative metabolite content are provided in the Supplementary files. The logarithmic transformation was applied to the metabolite abundance values as the majority of the metabolites followed a right-skewed distribution. They are further corrected for run and experimental batch effects by treating them as random, separately for the control and HNT conditions. A total of 66 metabolite abundance values were used as response variables for the subsequent genetic analysis. The list of metabolite contents analyzed in the current study is presented in Table S1.

### Genomic heritability of metabolite accumulation

A total of 162 rice lines with metabolite accumulation and SNP marker data in both control and HNT conditions were used for quantitative genetic analysis. The amount of metabolite content variability explained by SNP was evaluated by estimating genomic heritability. A single-trait genomic best linear unbiased prediction (GBLUP) model was

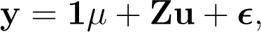

where **y** is a vector of metabolic abundance values; **1** is a vector of ones; *µ* is the overall mean; **Z** is the incidence matrix relating gene content to metabolic abundance values; **u** is a vector of random additive genetic values of the accessions; and ***ɛ*** is a vector of residuals. We assumed **u** *∼* N(**0**, 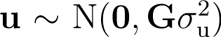), where 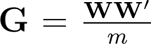 is the genomic relationship matrix between the accessions; 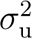 is the additive genetic variance; **W** is the centered and standardized gene content matrix; and *m* is the number of SNP markers. Further, we assumed 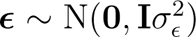, where **I** is the identity matrix and 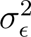 is the residual variance. Given the restricted maximum likelihood estimates of the variance components from the GBLUP model, genomic heritability estimates were calculated as 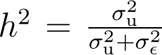. Genomic heritability estimates were inferred using the sommer R package (Covarrubias-Pazaran, 2016) for each metabolite accumulation individually.

### Single-trait genomic prediction of metabolites

The single-trait GBLUP model described above was used to evaluate the accuracy of genomic prediction of metabolite abundance values. Predictive performance was evaluated by repeated random subsampling cross-validation (CV), repeated 100 times. We split the accessions so that 80% and 20% of the accessions were included in the training and testing sets, respectively. The predictive performance of the model was assessed using the Pearson correlation between the observed metabolic abundance values and the predicted metabolic abundance values.

We also evaluated whether capturing non-additive genetic effects would result in improved prediction of metabolite abundance values using reproducing kernel Hilbert spaces regression (RKHS) (Gianola and Van Kaam, 2008). RKHS uses the Gaussian kernel (**GK**) instead of **G**. The Gaussian kernel matrix is equivalent to modeling additive by additive epistatic gene action up to infinite order by taking the Hadamard product between **G** matrices (Jiang and Reif, 2015). The **GK** is also known as a space continuous version of the diffusion kernel, which is deployed on graphs (Morota et al., 2013). The **GK** between a pair of accessions *i* and *j* with their SNP vectors **w***_i_ ∈* (0, 1, 2) and **w***_j_ ∈* (0, 1, 2) is given by:

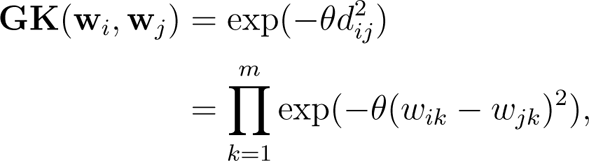

where 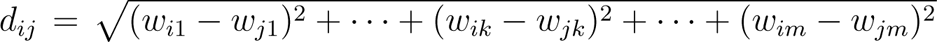 is the Euclidean distance and *θ* is the smoothing parameter. The role of *θ* is to smooth the relationship between the accessions. A large *θ* results in **GK** entries closer to 0 and a smaller *θ* results in entries closer to 1. Thus, *θ* controls the degree of genomic similarity between accessions. We applied kernel averaging (i.e., multiple kernel learning) by creating two **GK** matrices using extreme values of the smoothing parameters such that the means of the average off-diagonal elements of the corresponding kernels were 0.25 and 0.78, respectively.

The single-trait GBLUP and RKHS models were fit using the BGLR R package (Pérez and de los Campos, 2014). A flat prior was assigned to *µ*. The variance components, 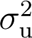 and 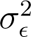, were drawn from a scaled inverse *χ*^2^ distribution with degrees of freedom *ν* = 5 and scale parameter *s* such that the prior means of the variance components equal half the phenotypic variance. A total of 50,000 Markov Chain Monte Carlo samples were used after 10,000 burn-ins with a thinning rate of 5 to obtain the posterior means for all unknowns.

### Genomic correlation between the same metabolite accumulation in different treatments

Genomic correlations between the same metabolite measured in control and HNT conditions were estimated by fitting bivariate GBLUP. A multi-trait model postulates phenotypic values of a genotype in different treatments (environments) as different but correlated traits (Hayes et al., 2016). Genetic correlations between treatments provide information about the relative magnitudes of genotypic values among accessions that change between treatments. A greater deviation from the genetic correlation of 1 indicates the presence of genotype by treatment (i.e., genotype by environment) interactions for metabolite accumulation. Genetic and residual variances in the single-trait GBLUP model were extended to the following variance-covariance structure.

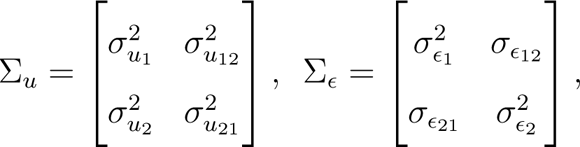

where subscripts 1 and 2 refer to the metabolite abundance values in the control and HNT conditions, respectively. In this parameterization, **u** and ***ɛ*** were assumed to follow a multivariate Gaussian distribution of **u** *∼ N* (0, Σ*_u_ ⊗* **G**) and *ɛ ∼ N* (0, Σ*_ɛ_ ⊗* **I**), respectively, where *⊗* is the Kronecker product. An inverse Wishart distribution was assigned to Σ*_u_* and Σ*_ɛ_* with degrees of freedom *ν* = 4 and scale matrix *S* such that the prior means of Σ*_u_* and Σ*_ɛ_* equal half of the phenotypic variance. The bivariate GBLUP was fit using 50,000 Markov chain Monte Carlo samples, 10,000 burn-ins, and a thinning rate of 5, implemented in the BGLR R package (Pérez-Rodríguez and de Los Campos, 2022). Since there were 66 metabolites, the above bivariate model was fitted 66 times.

### Exploratory factor analysis

To facilitate interpretation of the genomic heritability, genomic correlation, and genomic prediction results, exploratory factor analysis (Momen et al., 2021) was performed to identify underlying latent factors controlling *t* = 66 metabolites. All metabolites were modeled as

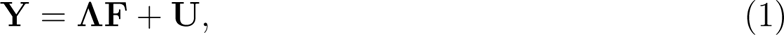

where **Y** is the *t × n* matrix of metabolic abundance values; **Λ** is the *t × q* matrix of factor loadings relating metabolite contents and latent common factors; **F** is the *q × n* matrix of latent factor scores; and **U** is the *t × n* vector of unique effects not explained by the *q* underlying latent common factors. Here, *n* is the number of accessions, *q* is the number of latent factors, and *t* is the number of metabolites. The variance-covariance matrix of **Y** was modeled as

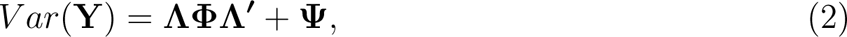

where *V ar*(**Y**) is the *t × t* variance-covariance matrix of the metabolites; **Φ** is the variance of the factor scores; and **Ψ** is a *t× t* diagonal matrix of unique variance. Here, the unknown parameters are **Λ**, **Φ**, and **Ψ**, which must be estimated from the data. We assumed **Φ** = **I**,which yields factors with unit variance (Jöreskog, 1967; Anderson, 2003). Assuming **F** *∼ N* (**0**, **I**), **Λ** and **Ψ** were estimated by maximizing the log-likelihood of *L*(**Λ**, **Ψ***|***Y**) coupled with the varimax rotation (Kaiser, 1958) using the psych R package (Revelle, 2018). A threshold of *λ > |*0.4*|* was used to screen out factor loadings, followed by assigning each metabolite to only one of the factors based on its largest loading. The optimal number of factors was determined by parallel analysis (Horn, 1965; Hayton et al., 2004). First, simulated data were generated from the observed data, and then the eigenvalues were extracted until the observed data had a smaller eigenvalue than the simulated data. The number of eigenvalues was considered as the number of optimal factors based on 20 simulated analyses.

### Simultaneous regression modeling of metabolites

Evaluation of the extent of correlations between different metabolite accumulation at the genetic level and the gain in genomic prediction by leveraging correlations between metabolites are of interest in quantitative genetics. Inference of genetic correlations between different metabolites within the same treatment and genomic prediction of metabolites considering all metabolites were performed separately in the control and HNT conditions using the MegaLLM R package (Runcie et al., 2021). MegaLLM is a multi-trait mixed effect factor model that exploits the factorization of genetic covariances between traits to speed up computation. The MegaLLM model was

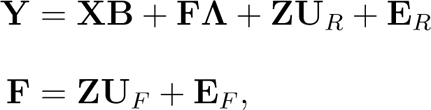

where **X** is the design matrix including the intercepts for each trait with the effect size matrix **B**; **F** is an *n× q* matrix of latent factors; **Λ** is a *q × t* factor loading matrix; and **U***_R_* and **U***_F_* are coefficients matrices. The following distributions were specified for the random effects.

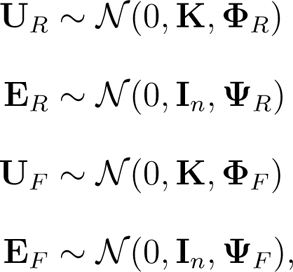

where *N* is the matrix normal distribution with mean, covariance between rows, and covariance between columns; **K** is a kernel matrix between accessions; **Φ***_R_* and **Ψ***_R_* are diagonal matrices of dimension *t*; and **Φ***_F_* and **Ψ***_F_* are diagonal matrices of dimension *q*. A total of 10,000 Markov Chain Monte Carlo samples after 6,000 burn-ins and 7,000 Markov Chain Monte Carlo samples after 4,000 burn-ins were used for genetic correlation and multi-trait genomic prediction analyses, respectively, with a thinning rate of 5 to obtain the posterior means for all unknowns. For MegaLMM-based genomic prediction of metabolite abundance values, we evaluated the importance of additive genetic variation and non-additive genetic variation by setting **K** = **G** (MegaLMM-G) and **K** = **GK** (MegaLMM-GK). The factorization implemented in MegaLMM allowed simultaneous fitting of all 66 metabolites to estimate their genetic correlations and perform multi-trait genomic prediction. The same CV scheme described in the single-trait genomic prediction was used.

### Deep learning models

Deep learning (DL), specifically convolutional neural networks (CNN), has proven effective for analyzing image data (Goodfellow et al., 2016). It has the ability to approximate any function given enough data to learn from. However, previous studies applying CNN in the context of genomic prediction have directly used a SNP marker matrix, **G** matrix, or its Cholesky decomposition as input predictors, which may not fully exploit the strength of CNN (Montesinos-Ĺopez et al., 2021). To take advantage of CNN capabilities, we transformed SNP data into images following the modified version of the DeepInsight framework (Sharma et al., 2019) and neural network modules. Briefly, the framework consists of rearranging similar SNPs into clusters, extracting image features, and fitting them into a regression model. An overview of these three steps is summarized in Figure 1. The first step transposes the original SNP matrix and splits it by chromosomes to form 12 sub-SNP matrices, then transforms each sub-SNP matrix into a feature matrix in the 2D Cartesian plane by taking the top two components from the t-SNE (Van der Maaten and Hinton, 2008), where a point represents a SNP marker. Here, a nonlinear dimensional reduction technique, t-SNE, was used to embed high-dimensional SNP data into a low-dimensional space of two dimensions. When two SNP markers are in high linkage disequilibrium, they are expected to have similar Cartesian coordinates. The convex hull algorithm was then applied to find the smallest rectangle containing all points and rotate it since CNN assumes that the input images are arranged in a horizontal or vertical form. This serves as a conversion from Cartesian coordinates to pixels. The pixel frame size is fixed to 277 *×* 277. Finally, the SNP codes (0 and 2) of each genotype were mapped to these pixel locations representing different shades of gray; 0 was represented by a lighter color, while 2 was depicted in a darker color. Chromosome-specific images were created for each genotype to capture small differences in SNP codes. These steps to transform tabular SNP data into images allow the full capabilities of CNN to be applied to genomic predictions. Since each genotype had 12 images corresponding to 12 chromosomes, we used multi-branch CNN regression. In particular, we considered the following CNN architectures: VGG16 (Simonyan and Zisserman, 2014), ResNet50 (He et al., 2016), EfficientNetB7 (Tan and Le, 2019), InceptionV3 (Szegedy et al., 2016), MobileNetV2 (Sandler et al., 2018), and DenseNet201 (Huang et al., 2017). Briefly, VGG16 emphasizes simplicity with deep convolutional layers, ResNet50 introduces skip connections for efficient training of deep networks, EfficientNetB7 uses balanced scaling of network dimensions, InceptionV3 uses inception modules for diverse receptive fields, MobileNetV2 focuses on lightweight architecture with inverted residuals, and DenseNet201 maximizes feature reuse through densely connected blocks. Due to the relatively small number of genotypes in our dataset, which typically requires thousands of images, we took advantage of transfer learning. Specifically, we used pre-training weights for the layers (excluding the top layers) from CNN models trained on the ImageNet dataset (Deng et al., 2009). Applying these weights to each SNP image facilitated feature extraction, with the same weights shared across 12 branches. We then used the optimal ridge regression model, selected from regularization strengths of 10, 100, 1,000, and 10,000, to predict metabolite abundance values. The same CV scheme described for single-trait genomic predictions was used. We implemented these models using Python version 3.9.0 and the deep learning framework TensorFlow version 2.12.0 with the Keras API version 2.6.0.

**Figure 1:**
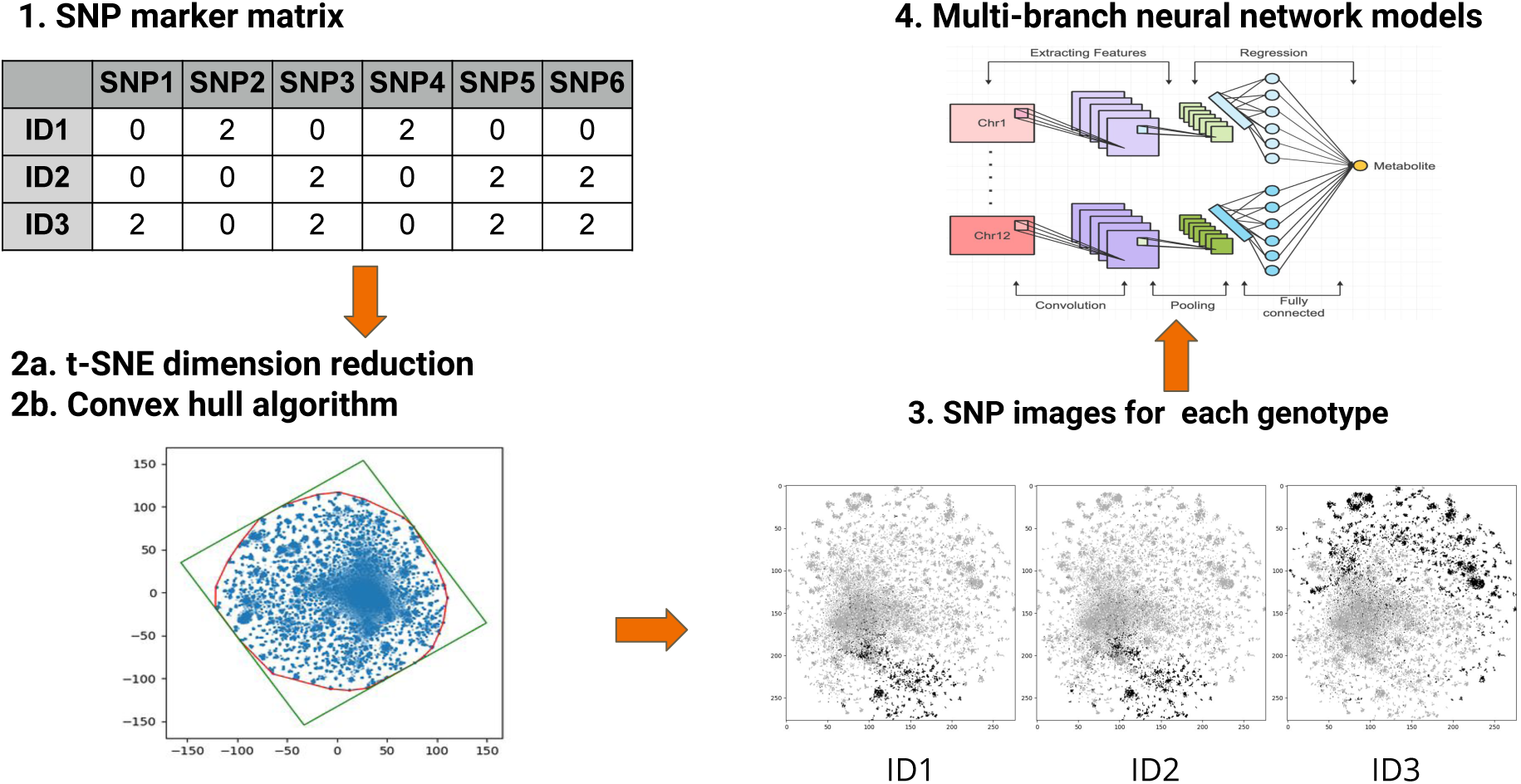
Flowchart of converting single nucleotide polymorphisms to image data.

## Results

### Single-trait analysis of metabolites

Pairwise Pearson correlations between metabolic profiles are shown in Figure S1. Pairs between leucine-valine, isoleucine-valine, isoleucine-leucine, asparagine-dihydrouracil, pantothenic acid-glyceric acid, ornithine-citrulline, and adenosine-uridine in the control condition, as well as leucine-valine, isoleucine-valine, isoleucine-leucine, asparagine-dihydrouracil, adenine-glyceric acid, and 4-hydroxy-3-methoxybenzoic acid-hydroxybenzoic acid in the HNT condition showed positive correlation coefficients greater than 0.8. Genomic heritability estimates for metabolite accumulation are shown in Figure 2. The majority of metabolite abundance values were low to moderately heritable (Figure 2A). Genomic heritability estimates ranged from 0 to 0.80 and 0 to 0.70 in control and HNT conditions, respectively. Overall, metabolite accumulation showed larger genomic heritability estimates in control compared to HNT conditions (Figure 2B). Their mean (median) values were 0.27 (0.24) and 0.22 (0.19) in control and HNT conditions, respectively. The Pearson correlation coefficient of genomic heritability estimates between control and HNT conditions was 0.50, indicating that the two treatments have a modestly similar pattern. Although most metabolites showed agreement of heritability estimates between control and HNT conditions, some metabolites deviated from this trend (Figure 2C). A total of 24 and 29 metabolites, respectively, showed small (*<* 0.05) and large (*>* 0.1) heritability differences between control and HNT conditions. Genomic correlation estimates of the same metabolites between the two treatments are shown in Figure 3. All metabolites showed genomic correlation estimates less than 1, indicating genotype by environment interactions to some extent (Figure 3A). The genomic correlation estimates ranged from 0.12 to 0.86. Their mean and median values were 0.48 and 0.49, respectively (Figure 3B).

**Figure 2:**
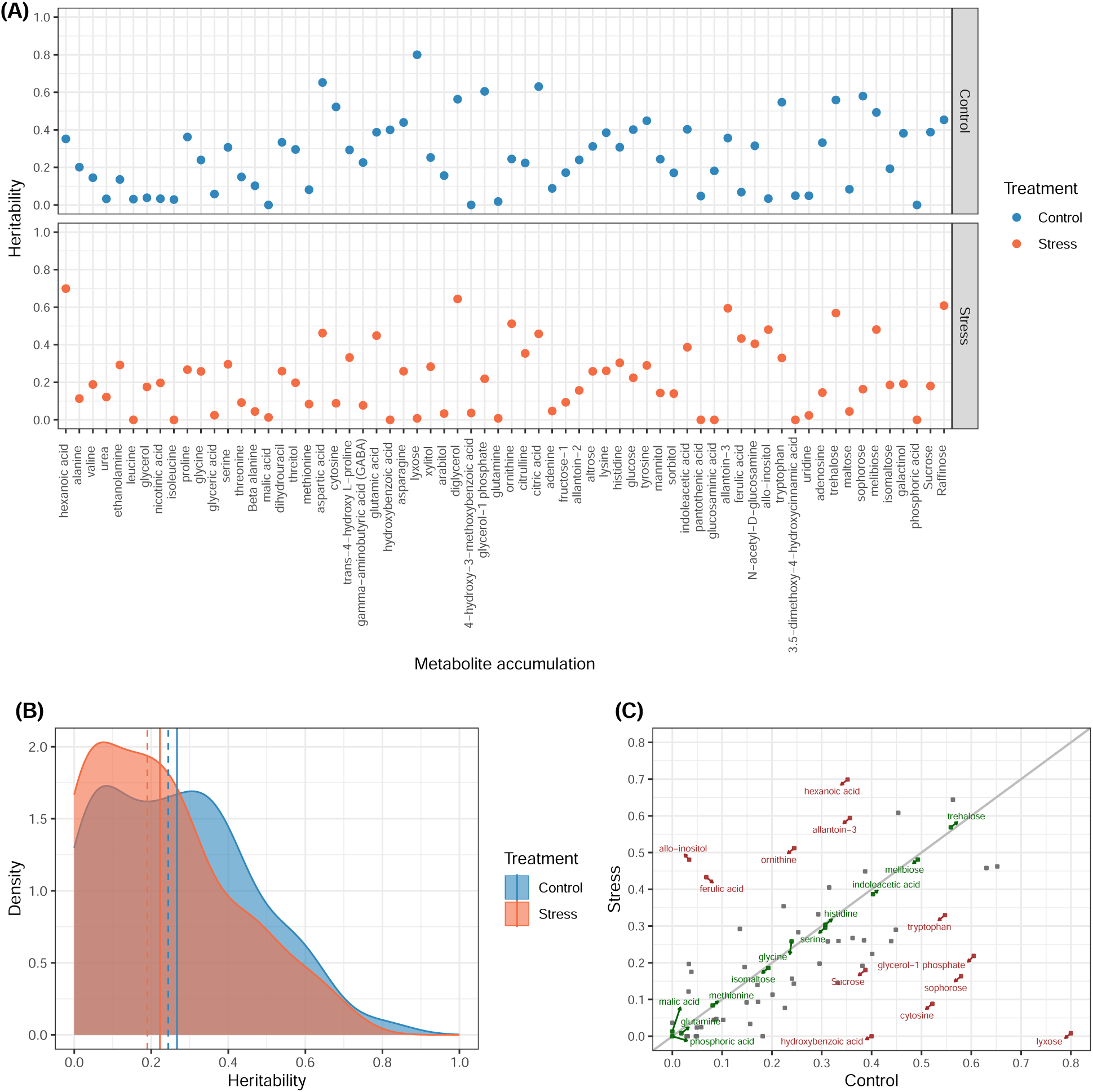
Genomic heritability estimates of metabolite accumulation in control and high night temperature stress conditions. A) Scatter plot. B) Density plot. Solid and dashed lines indicate mean and median, respectively. C) Agreement of heritability estimates between control and high night temperature stress conditions. Metabolite accumulation in green and red colors indicate that the heritability difference between control and high night temperature stress conditions was small (*<* 0.05) and large (*>* 0.1).

**Figure 3:**
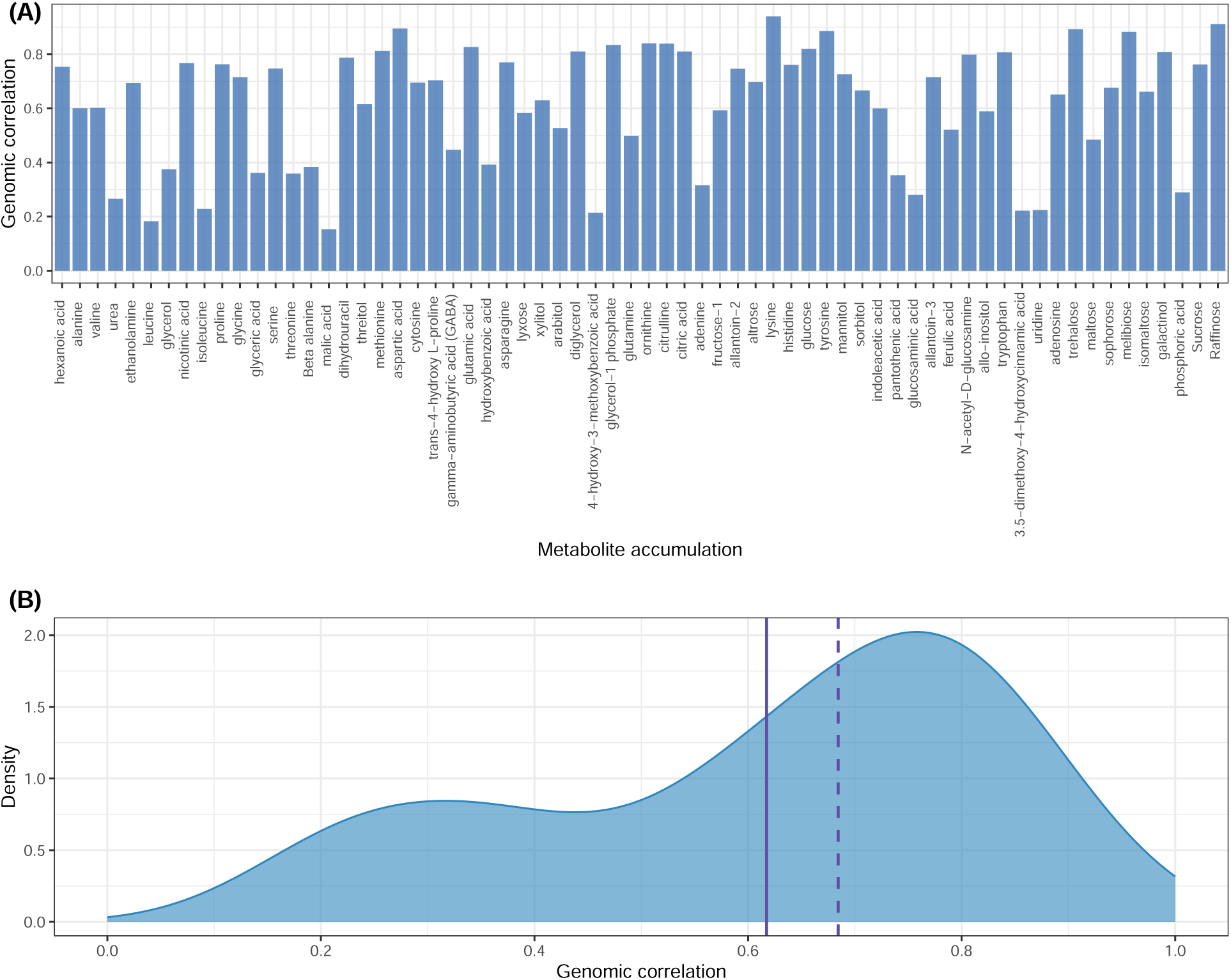
Genomic correlation estimates between the same metabolite accumulation measured under control and high night temperature stress conditions. A) Scatter plot. B) Bar chart. Solid and dashed lines indicate mean and median, respectively.

Genomic prediction accuracy of metabolite accumulation in control and HNT conditions is shown in Figure 4. The results varied widely between metabolites (Figure 4A). The accuracy of genomic predictions ranged from 0.01 to 0.66 and 0.01 to 0.63 in control and HNT conditions, respectively. Consistent with genomic heritability estimates, genomic prediction accuracies were higher in control compared to HNT conditions (Figure 4B). Their mean (median) values were 0.32 (0.32) and 0.28 (0.29) in control and HNT conditions, respectively. Similar to the genomic heritability estimates obtained, although most metabolites showed an agreement of genomic prediction accuracy between control and HNT conditions, there were some metabolites that deviated from this trend (Figure 4C). A total of 24 and 20 metabolites, respectively, showed small (*<* 0.05) and large (*>* 0.1) differences in genomic prediction accuracy between control and HNT conditions.

**Figure 4:**
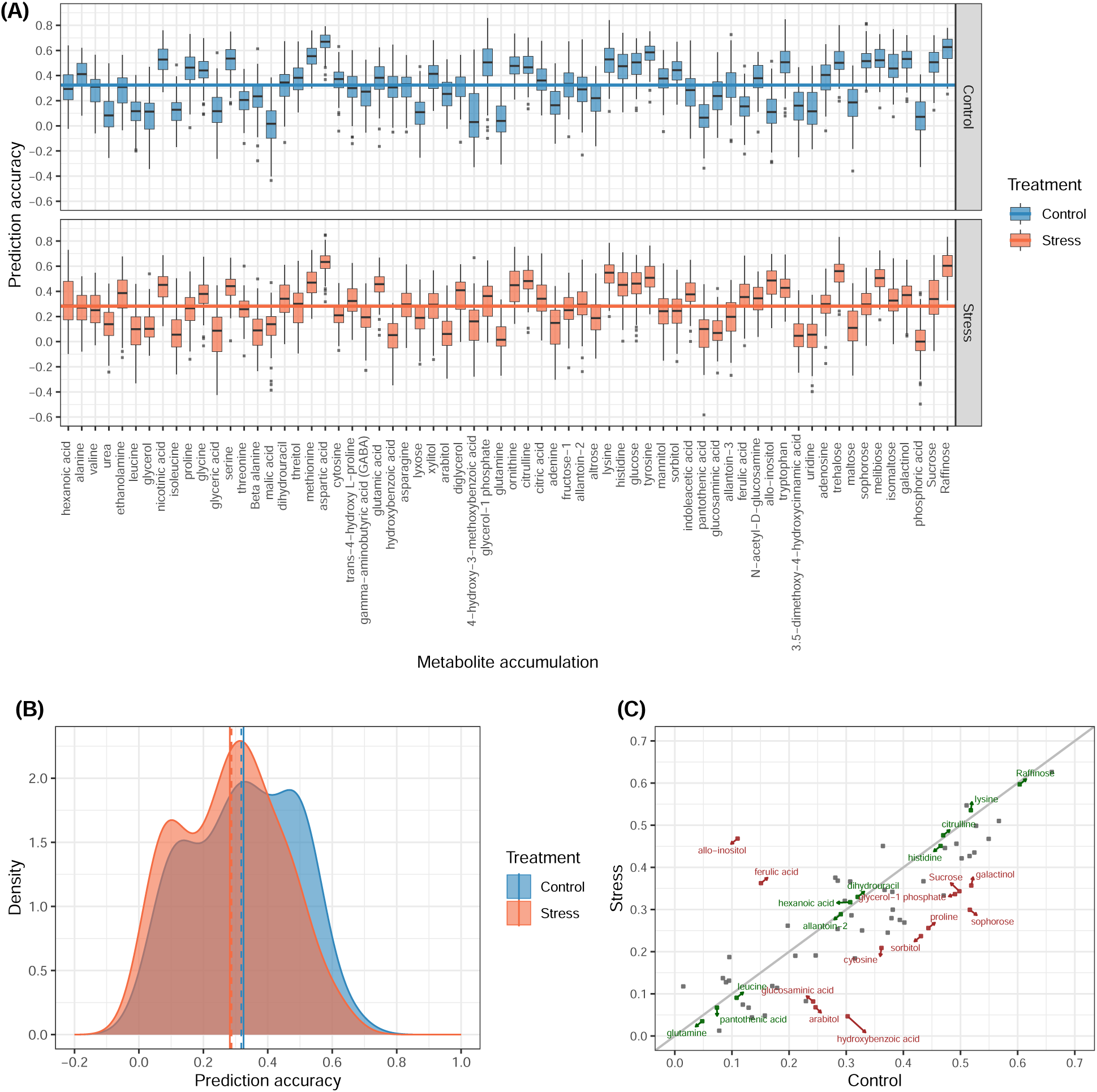
Genomic prediction accuracy of metabolite accumulation in control and high night temperature stress conditions. A) Box plot. The horizontal line indicates the mean value. B) Density plot. The solid and dashed lines indicate the mean and median, respectively. C) Agreement of genomic prediction accuracy between control and high night temperature stress conditions. Metabolite accumulation in green and red colors indicate that the genomic prediction difference between control and high night temperature stress conditions was small (*<* 0.05) and large (*>* 0.1).

There was also a linear relationship between genomic heritability estimates and genomic prediction accuracies. The coefficients of determination obtained from ordinary least squares between genomic heritability estimates and genomic prediction accuracies squared were 0.89 and 0.88 for the control and HNT conditions, respectively.

### Latent factors behind metabolites

Parallel analysis was used to find the optimal number of latent factors. In both the control and HNT conditions, the first five eigenvalues from the real data were larger than the first five eigenvalues from the simulated random data. Therefore, five latent factors were investigated in the next step. Exploratory factor analysis was performed to understand the biological significance of the five latent factors by examining the covariation among the measured metabolite accumulation. The magnitude of the contributions of the latent factors to metabolite accumulation in the control and HNT conditions is summarized in Figure S2. Because explanatory factor analysis allows for cross-loading of metabolites, the largest loading for each metabolite accumulation was selected based on a threshold of *> |*0.4*|*. This resulted in each metabolite loading on only one factor, except that urea, glycine, methionine, arabitol, diglycerol, citric acid, allo-inositol, trehalose, maltose, and sophorose, and hexanoic acid, urea, threitol, lyxose, arabitol, diglycerol, and citric acid did not load on any factors in the control and HNT conditions, respectively.

Density plots of genomic heritability, genomic correlation, and genomic prediction accuracy categorized into 10 latent factors are shown in Figure S3. Here, the first five (F1-F5) and last five (F6-F10) latent factors denote factors controlling the control and HNT conditions, respectively. In the control condition, F2 (0.43) and F4 (0.30) had higher mean estimates of genomic heritability, and F2 (0.51), F5 (0.39), and F4 (0.27) had higher means of genomic prediction accuracy. In the HNT condition, F10 (0.33), F7 (0.32), and F8 (0.30) had higher mean estimates of genomic heritability, and F7 (0.45), F8 (0.28), and F10 (0.27) had higher means of genomic prediction accuracy.

### Single-trait analysis using non-additive models

The gain in prediction accuracy provided by single-trait RKHS or DL-based on different architectures relative to single-trait GBLUP is shown in Figure 5. The inclusion of epistasis increased the prediction accuracy of some metabolite accumulation in RKHS. For example, a gain in prediction was observed for allo-inositol, glyceric acid, uridine, trans-4-hydroxy L-proline, and allantoin-2 in the control conditions as well as for urea, hexanoic acid, glycerol, maltose, allantoin-2, and N-acetyl-D-glucosamine in the HNT condition. Capturing epistasis using RKHS resulted in an average gain of 0.34% and -2.78% for the control and HNT conditions, respectively. Overall, the use of DL by converting SNP into image data increased prediction accuracy more than RKHS. An example of a set of SNP transformed into image data for a randomly selected genotype is shown in Figure 6. EfficientNetB7, ResNet50, and VGG16 showed noticeable gains in the control conditions. In particular, in the control conditions, VGG16, EfficientNetB7, and ResNet50 improved prediction accuracy in 30, 26, and 8 metabolite contents, respectively. In the HNT conditions, VGG16 and MobileNetV2 showed a gain in prediction accuracy in 23 and 13 metabolite contents, respectively.

**Figure 5:**
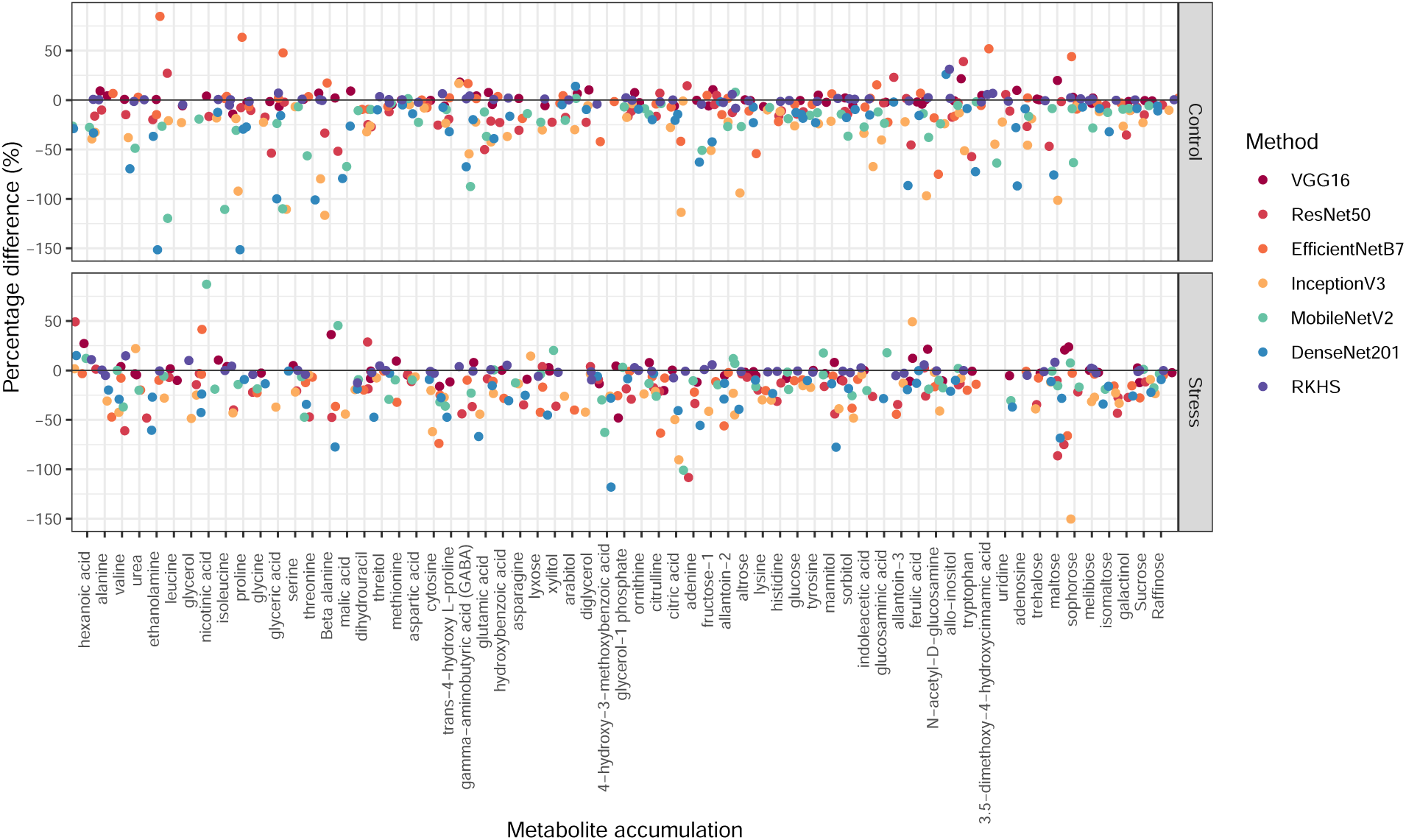
Percentage difference of gain in prediction accuracy for single-trait reproducing kernel Hilbert spaces regression (RKHS), VGG16, ResNet50, EfficientNetB7, InceptionV3, MobileNetV2, and DenseNet201 relative to single-trait genomic best linear unbiased prediction.

**Figure 6:**
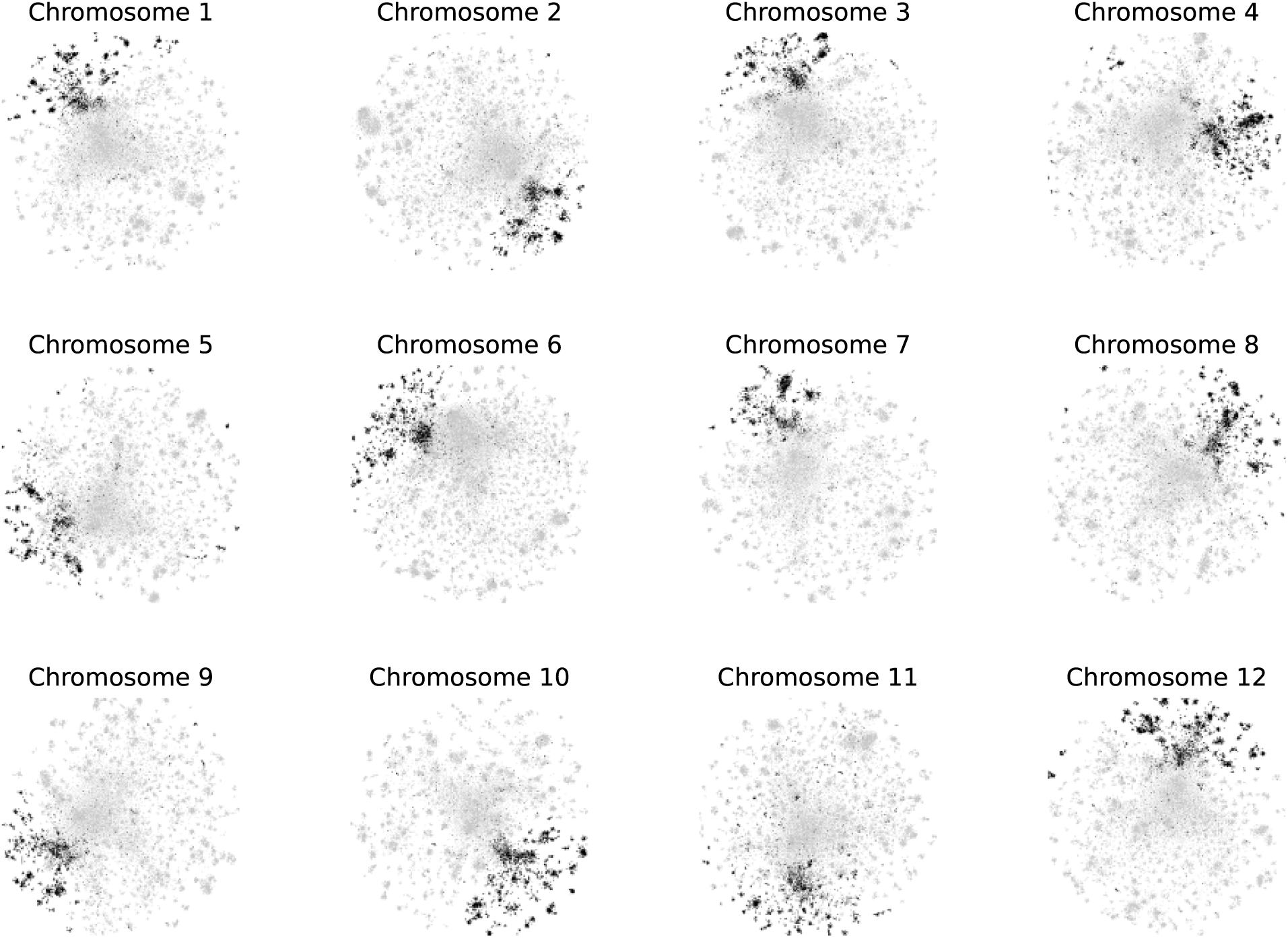
Example of a set of single nucleotide polymorphisms transformed into image data for a randomly selected genotype. Images of 12 chromosomes were processed in the multi-channel convolutional neural networks.

### Joint analysis of multiple metabolites

Simultaneous regression modeling of all metabolite abundance values was performed using MegaLMM. Estimates of genomic correlations between different metabolite abundance values are shown in Figure S4. Estimates ranged from -0.28 to 0.79 and -0.20 to 0.74 for the control and HNT conditions, respectively. Similar to the results for genomic heritability and single-trait genomic prediction, genomic correlation estimates were larger in the control than in the HNT conditions (Figure 7). Their mean (median) values were 0.14 (0.17) and 0.12 (0.16) in the control and HNT conditions, respectively.

**Figure 7:**
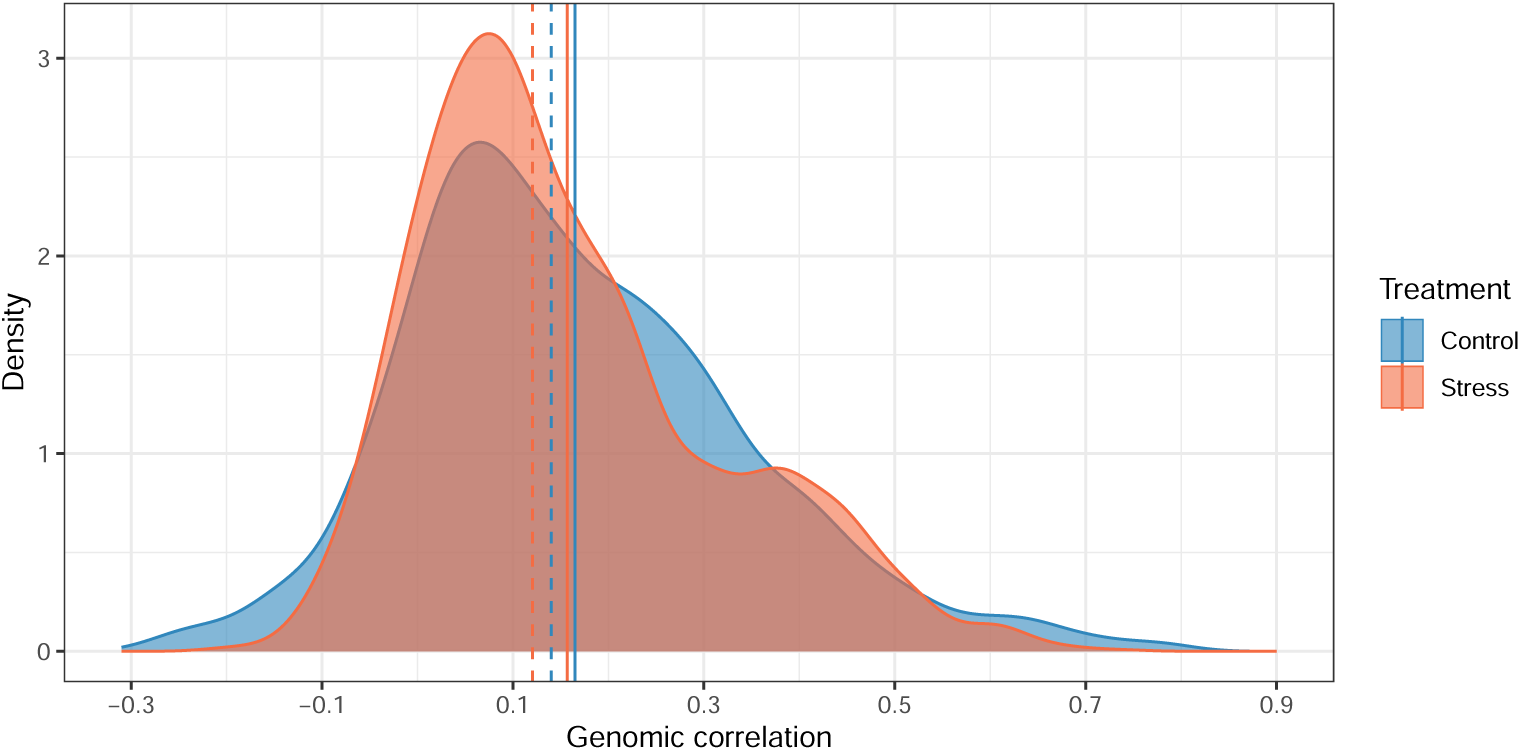
Genomic correlation estimates between different metabolite accumulation in control and high night temperature stress conditions. The solid and dashed lines indicate mean and median, respectively.

The gain in prediction accuracy provided by multi-trait MegaLMM-G and multi-trait MegaLMM-GK relative to single-trait GBLUP is shown in Figure 8. Using the correlation between metabolites through simultaneous fitting increased the genomic prediction accuracy of some metabolites, including threonine, gamma-aminobutyric acid, allo-inositol, uridine, and maltose in the control condition as well as proline, malic acid, xylitol, 4-hydroxy-3-methoxybenzoic acid, and sucrose in the HNT condition. When the total metabolites were fitted together with **G** using MegaLMM-G, the average gain in prediction accuracy was +0.48% and -0.45% for the control and HNT conditions, respectively (Figure 8B). Finally, combining MegaLMM with **GK** (MegaLMM-GK) resulted in the largest average gain of 4.80% and 1.79% for the control and HNT conditions, respectively (Figure 8C). Overall, the use of MegaLMM was not successful for all metabolites, as some metabolites showed lower prediction accuracies than those of GBLUP.

**Figure 8:**
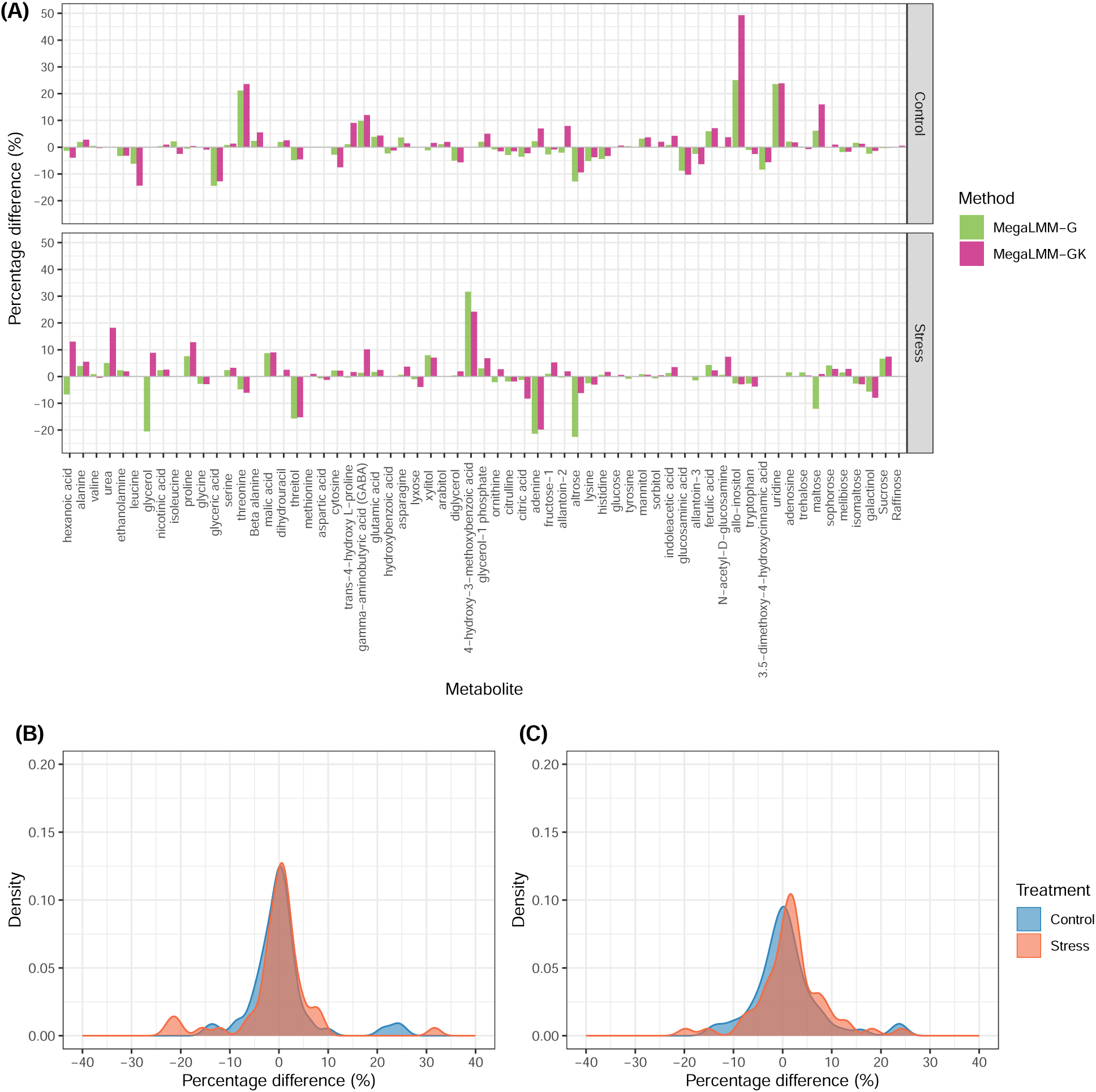
Percentage difference of gain in prediction accuracy for multi-trait genomic best linear unbiased prediction (MegaLMM-G) and multi-trait reproducing kernel Hilbert spaces regression (MegaLMM-GK) relative to single-trait genomic best linear unbiased prediction (A). Density plots of percentage difference are shown for MegaLMM-G (B) and MegaLMMGK (C).

## Discussion

In rice, HNT stress negatively affects grain yield and quality. It has been reported that an increase in 1*^◦^*C average minimum temperature during the growing season results in a 10% yield loss (Peng et al., 2004). Therefore, there is a need to better understand the response of rice to HNT stress and to improve the resilience of rice to HNT stress. Studies of metabolites as phenotypes using a quantitative genetic approach via whole-genome data have recently increased in plants (Guo et al., 2020; Campbell et al., 2021; Brzozowski et al., 2022; Onogi, 2023). Although metabolic profiles of plants can be used to measure their HNT responses (Obata and Fernie, 2012), the use of whole-genome regression analysis to analyze metabolite accumulation has been limited in rice. In this study, we 1) quantified the extent of genetic control of variation in metabolite levels and 2) evaluated the accuracy of genomic prediction of metabolite accumulation under control and HNT conditions.

Overall, our study revealed low to moderate genomic heritability estimates for the metabolic profiles of rice accessions, consistent with previous findings of higher broad-sense heritability for secondary metabolites compared to primary metabolites in rice grains as assessed using line replications (Matsuda et al., 2012). Our results show that the observed pattern of low to moderate heritability extends to genome-based heritability estimates. Although a genomewide association study was beyond the scope of this study, previous research suggests that secondary metabolites tend to be controlled by a limited number of loci with large effects, whereas primary metabolites tend to be controlled by loci with smaller effects (Chen et al., 2014).

The accuracy of genomic prediction results provided by GBLUP for metabolite accumulation was consistent with the upper bound established by genomic heritability estimates. This observation was supported by a linear relationship between genomic heritability estimates and squared genomic prediction accuracies. The control conditions yielded higher genomic heritability estimates and genomic prediction accuracies compared to the HNT conditions, suggesting that accurate estimation of SNP effects was more difficult under the HNT condition or the presence of differential genetic control of metabolite accumulation between the two treatments. Several metabolites showed notable agreement or difference in genomic heritability estimates or genomic prediction accuracies between the control and HNT conditions. For example, allo-inositol and ferulic acid showed differences between the two conditions in both genomic heritability and genomic prediction analyses. These two metabolites are often related, although they are not related in metabolic pathways. They were more heritable and predictable under HNT than under control conditions. Ornithine and allantoin-3 were also found to differ between the two conditions in genomic heritability estimates related to nitrogen metabolism. Sucrose, proline, sorbitol, and galactinol showed higher prediction accuracy in genomic predictions under the control condition. Overall, amino acids tended to be enriched in both conditions.

The degree of genotype by environment interactions was assessed using a bivariate model in which the traits measured in the different environments (i.e., treatments) were treated as distinct but correlated traits. Estimates of genetic correlation between the same trait in different environments obtained from the multi-trait model reflect the degree of re-ranking of genotypes between environments. In general, genetic correlations of less than 0.8 are usually considered to represent different environments (Robertson, 1959). Since many of the genetic correlation estimates of the metabolite were less than 0.8, the extent of genotype by environment interactions in rice grain metabolites is not weak and indicates that re-ranking among accessions occurs between the control and HNT conditions.

To facilitate interpretation of the large number of metabolites analyzed in this study, we used exploratory factor analysis. Exploratory factor analysis is a statistical technique that can reveal the underlying structure among phenotypes when no prior assumptions can be made about the nature of the underlying factor. Factors F3 and F9 contained branched-chain amino acids, or BCAA, and factors F4 and F8 contained N-rich compounds. Of these, F4 and F8 were identified as having higher mean estimates of genomic heritability and mean accuracies of genomic prediction in the control and HNT conditions, respectively, in single-trait metabolite analysis.

The comparison between GBLUP and RKHS or DL suggests that the prediction accuracy of metabolite accumulation can be improved by capturing non-additive genetic effects for some, but not all, metabolites. In particular, the prediction gain provided by DL was remarkable for some metabolites, suggesting that transforming SNP data into image data is a potentially useful approach for genomic prediction by improving feature and structure extraction from non-image data. Our results are consistent with a recent study that reported encouraging results using the original version of DeepInsight for genomic prediction in maize (Galli et al., 2022). The modified DL approach used in the current study follows that of the original version in the image transformation step (Sharma et al., 2019), but differs in the adapted CNN architectures. It takes advantage of the recent development of CNN architectures coupled with multi-channel CNN. Furthermore, our results indicate that subsets of rice grain metabolite contents are controlled by non-linear genetic effects. However, these metabolites were not enriched in specific metabolic categories. Overall, the use of RKHS or DL was not useful for all metabolites because their prediction accuracies were lower than those of GBLUP.

Using MegaLMM allowed us to fit all metabolite accumulation simultaneously, taking advantage of the correlations between them. Otherwise, assuming a bivariate analysis was performed, (66 *×*65) / 2 = 2,145 pairs of models had to be fitted, which is computationally demanding. Joint quantitative genetic analysis of multiple metabolite accumulation, including genomic correlation and genomic prediction analyses, was provided by MegaLMM. Consistent with the single-trait metabolite analysis, the control conditions yielded higher genomic correlation estimates between metabolites and greater genomic prediction accuracies compared to the HNT conditions when MegaLMM was used. This further supports the hypothesis that accurate estimation of SNP effects was more difficult in the HNT condition or that there was differential genetic control of metabolites between the two treatments. Across all metabolites, MegaLMM-GK provided the best prediction on average compared to single-trait GBLUP. This result suggests that incorporating the correlation between metabolite accumulation and epistasis was an important factor in improving genomic predictions of metabolite accumulation.

## Conclusions

Primary metabolite accumulation from rice grains was low to moderately heritable. Genomic heritability estimates were slightly higher in control than in HNT. Genomic prediction accuracies of metabolite accumulation obtained from GBLUP were within the expected upper limit set by their genomic heritability estimates. Genomic correlation estimates for the same metabolite between the control and HNT conditions deviated from 1, indicating the presence of genotype by environment interactions. There was variation among metabolites in how well they agreed between the control and HNT conditions in terms of genomic heritability estimates and genomic prediction accuracies. Genomic prediction models such as RKHS and image-based DL for single-trait metabolite analysis were effective in improving prediction accuracy by capturing non-additive genetic effects for some, but not all, metabolites. Joint analysis of many metabolites via MegaLMM was also shown to be useful for improving genomic prediction accuracy by exploiting correlations among metabolite accumulation. The current study serves as a first important step to evaluate the genomic prediction accuracy of metabolites under control and HNT conditions.

## Author contributions

Ye Bi: Conceptualization; Formal analysis, Methodology; Visualization; Writing-original draft. Harkamal Walia: Investigation, Data curation, Funding acquisition; Writing-review & editing. Toshihiro Obata: Investigation, Data curation, Funding acquisition; Writing-review & editing. Gota Morota: Conceptualization; Methodology; Funding acquisition; Project administration; Supervision; Writing-original draft; Writing-review & editing.

## Data availability

Genotypic data regarding the rice accessions are available at the rice diversity panel website (http://www.ricediversity.org/). Scripts used in this work are publicly available in GitHub (https://github.com/yebigithub/GBLUP4Met).

## Abbreviations

(CNN): convolutional neural networks
(CV): cross-validation
(DL): deep learning
(GK): Gaussian kernel
(G): genomic relationship matrix
(GBLUP): genomic best linear unbiased prediction
(HNT): high night temperature
(RKHS): reproducing kernel Hilbert spaces
(SNP): single nucleotide polymorphism

