## Supplementary for "Genomic prediction of metabolic content in rice grain in response to warmer night conditions"

### Tables

Table S1: List of metabolites analyzed in the current study.

| Number | Metabolite | Number | Metabolite |
|---|---|---|---|
| 1 | hexanoic acid | 2 | alanine |
| 3 | valine | 4 | urea |
| 5 | ethanolamine | 6 | leucine |
| 7 | glycerol | 8 | nicotinic acid |
| 9 | isoleucine | 10 | proline |
| 11 | glycine | 12 | glyceric acid |
| 13 | serine | 14 | threonine |
| 15 | Beta alanine | 16 | malic acid |
| 17 | dihydrouracil | 18 | threitol |
| 19 | methionine | 20 | aspartic acid |
| 21 | cytosine | 22 | trans-4-hydroxy L-proline |
| 23 | gamma-aminobutyric acid (GABA) | 24 | glutamic acid |
| 25 | hydroxybenzoic acid | 26 | asparagine |
| 27 | lyxose | 28 | xylitol |
| 29 | arabitol | 30 | diglycerol |
| 31 | 4-hydroxy-3-methoxybenzoic acid | 32 | glycerol-1 phosphate |
| 33 | glutamine | 34 | ornithine |
| 35 | citrulline | 36 | citric acid |
| 37 | adenine | 38 | fructose-1 |
| 39 | allantoin-2 | 40 | altrose |
| 41 | lysine | 42 | histidine |
| 43 | glucose | 44 | tyrosine |
| 45 | mannitol | 46 | sorbitol |
| 47 | indoleacetic acid | 48 | pantothenic acid |
| 49 | glucosaminic acid | 50 | allantoin-3 |
| 51 | ferulic acid | 52 | N-acetyl-D-glucosamine |
| 53 | allo-inositol | 54 | tryptophan |
| 55 | 3.5-dimethoxy-4-hydroxycinnamic acid | 56 | uridine |
| 57 | adenosine | 58 | trehalose |
| 59 | maltose | 60 | sophorose |
| 61 | melibiose | 62 | isomaltose |
| 63 | galactinol | 64 | phosphoric acid |
| 65 | Sucrose | 66 | Raffinose |

### Figures

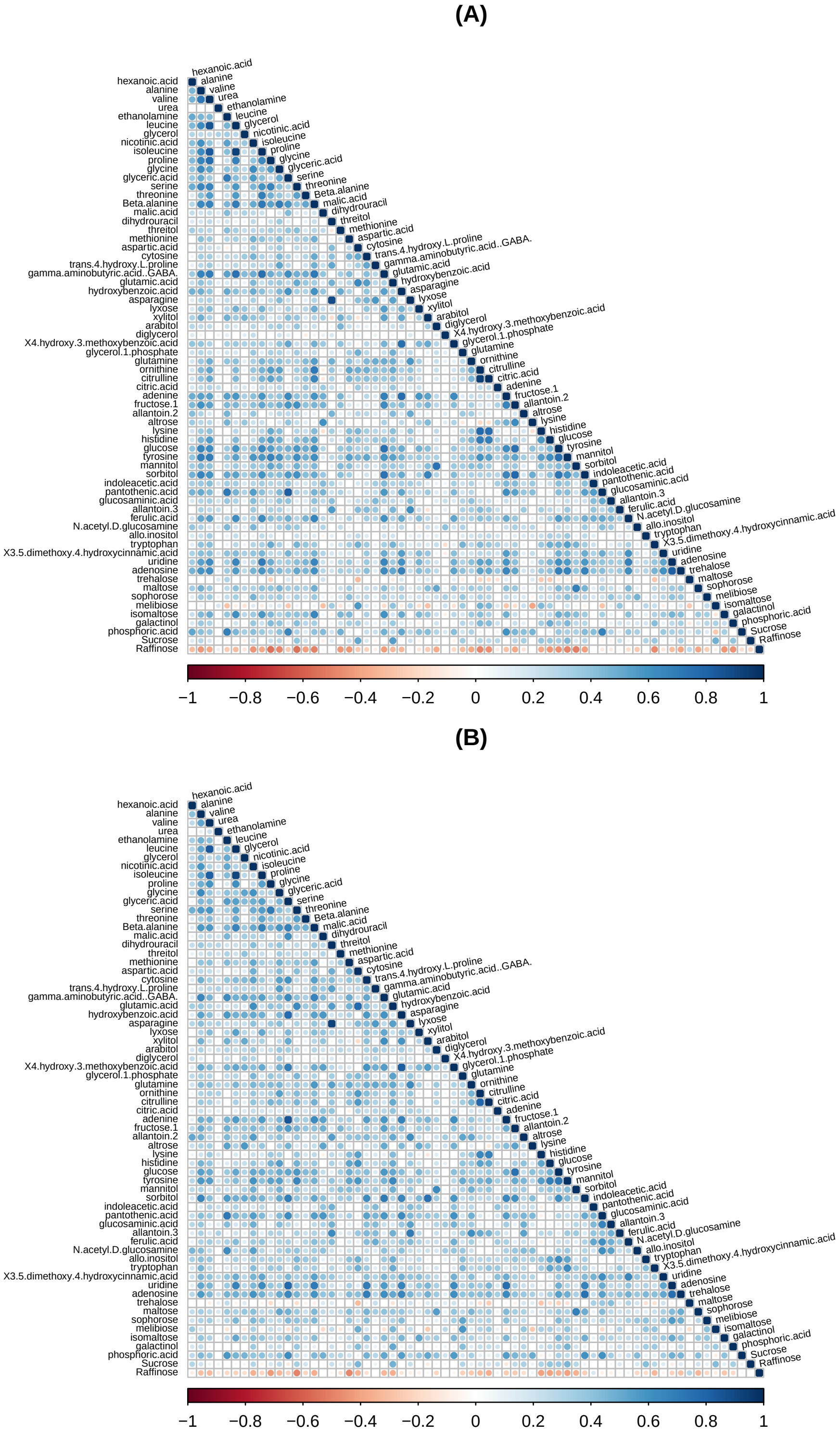


Figure S1: Pearson correlation heat maps between metabolic profiles in control (A) and high night temperature stress (B).

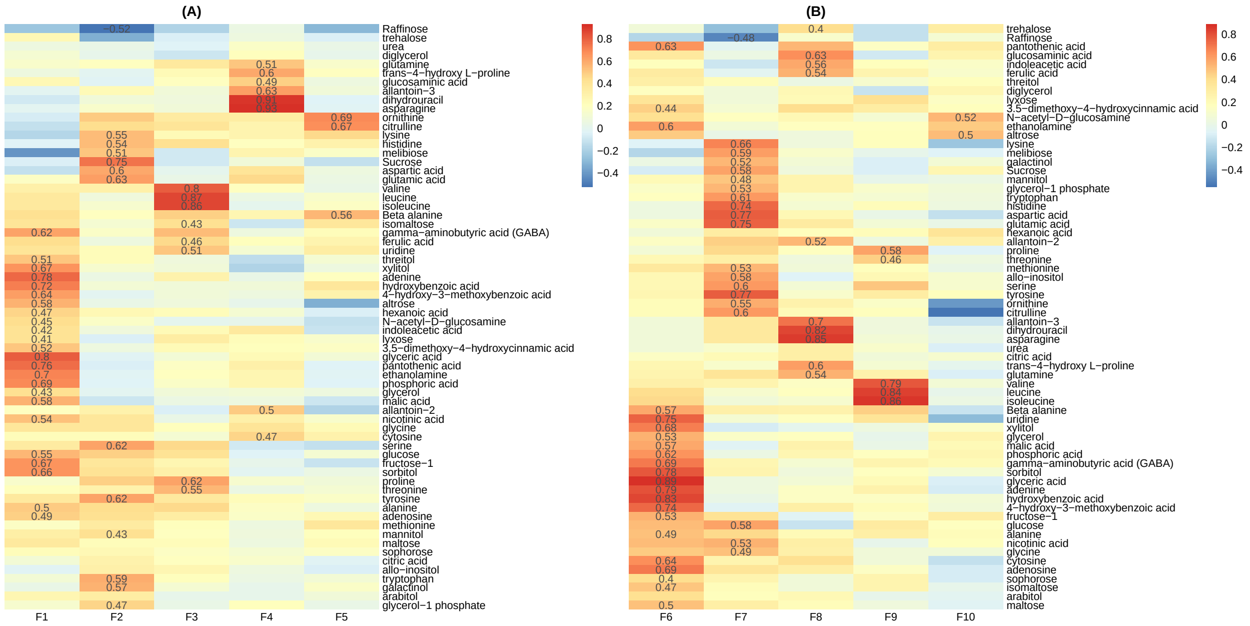


Figure S2: Heat maps of factor loadings after removing cross-loading by setting a cutoff of $\lambda > |0.4|$. The rows of each panel correspond to the observed metabolite accumulation and the columns correspond to the five latent factors in control (A) and high night temperature stress conditions (B).

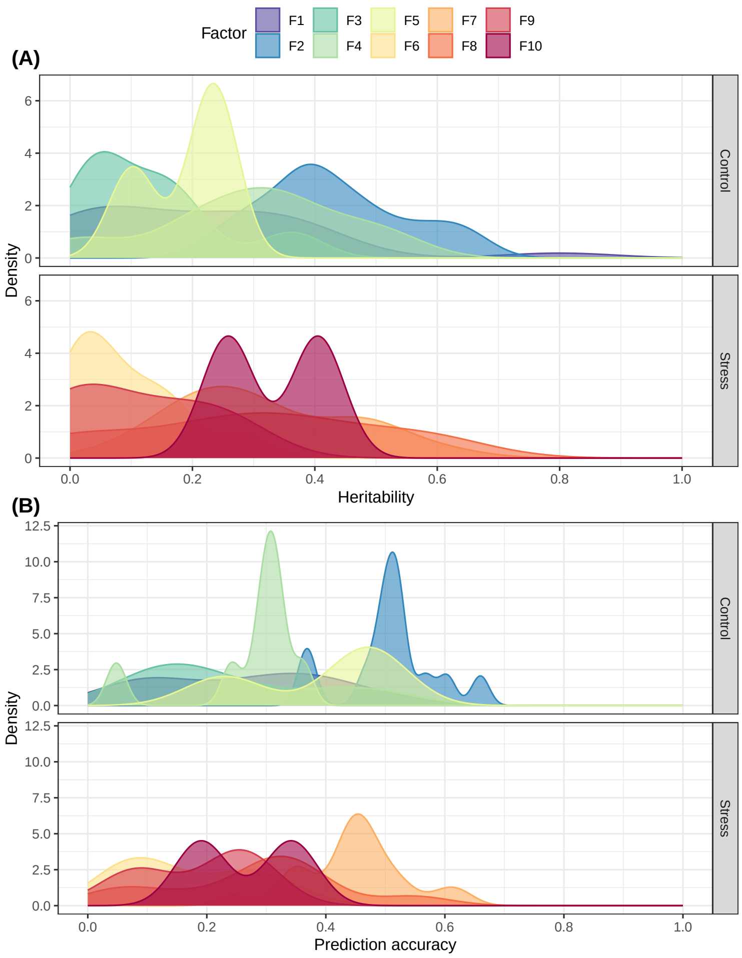


Figure S3: Genomic heritability estimates of metabolite contents in control and high night temperature stress conditions (A) and genomic prediction accuracy of metabolite contents in control and high night temperature stress conditions, separated by latent factors (B). F1-F5 and F6-F10 are latent factors in control and high night temperature stress conditions, respectively.

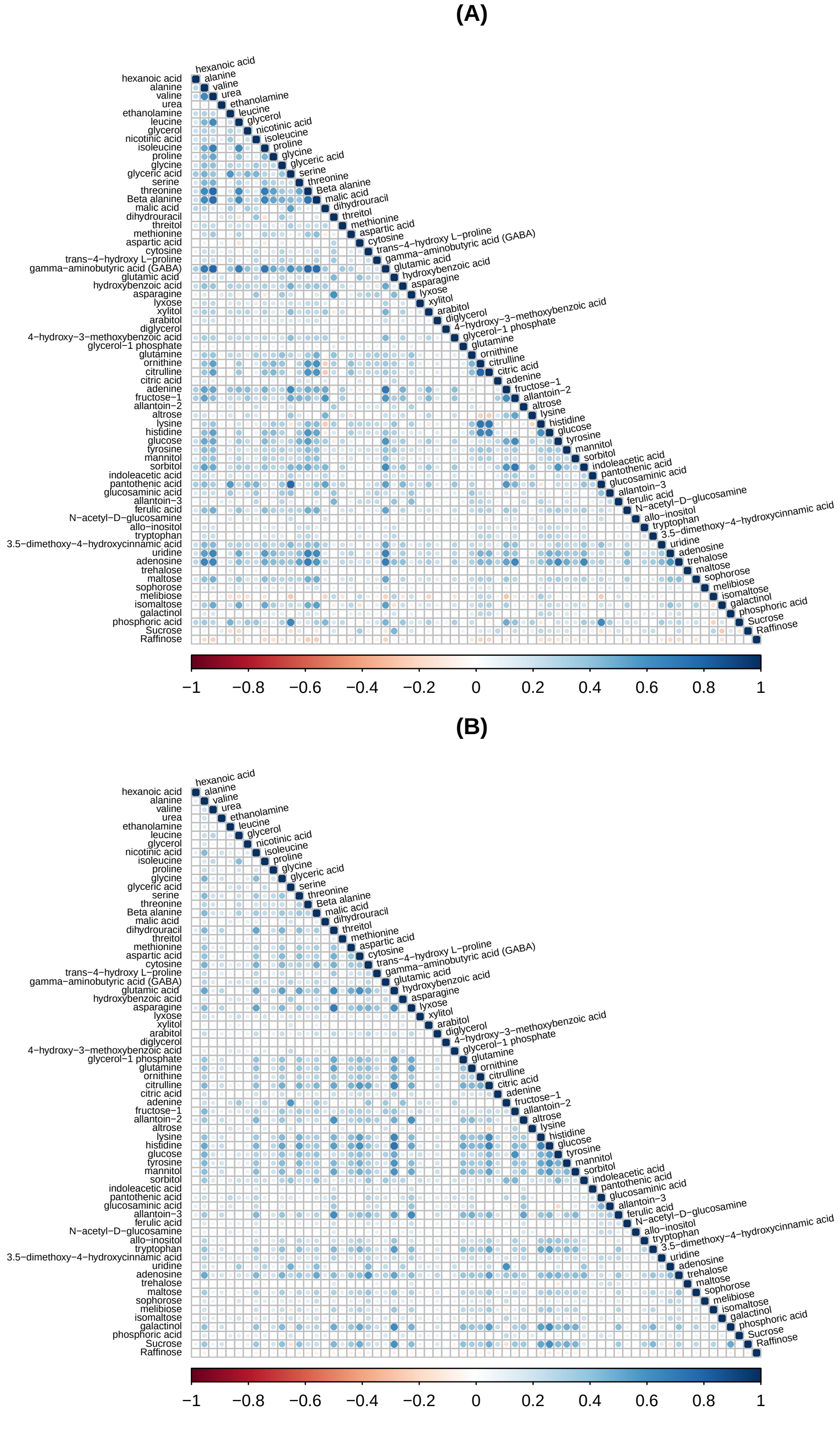


Figure S4: Genomic correlation estimates between different metabolites in control (A) and high night temperature stress (B).
